# TopicNet: a framework for measuring transcriptional regulatory network change

**DOI:** 10.1101/862177

**Authors:** Shaoke Lou, Tianxiao Li, Xiangmeng Kong, Jing Zhang, Jason Liu, Donghoon Lee, Mark Gerstein

## Abstract

Next generation sequencing data highlights comprehensive and dynamic changes in the human gene regulatory network. Moreover, changes in regulatory network connectivity (network “rewiring”) manifest different regulatory programs in multiple cellular states. However, due to the dense and noisy nature of the connectivity in regulatory networks, directly comparing the gains and losses of targets of key TFs is not that informative. Thus, here, we seek a abstracted lower-dimensional representation to understand the main features of network change. In particular, we propose a method called TopicNet that applies latent Dirichlet allocation (LDA) to extract meaningful functional topics for a collection of genes regulated by a TF. We then define a rewiring score to quantify the large-scale changes in the regulatory network in terms of topic change for a TF. Using this framework, we can pinpoint particular TFs that change greatly in network connectivity between different cellular states. This is particularly relevant in oncogenesis. Also, incorporating gene-expression data, we define a topic activity score that gives the degree that a topic is active in a particular cellular state. Furthermore, we show how activity differences can highlight differential survival in certain cancers.

## Introduction

In recent years, large-scale data on the interaction between proteins and genes has enabled the construction of complex transcriptional regulatory networks (Zhang et al., 2014, Liu et al., 2015). These networks model the molecular program for gene transcription by representing genes and regulatory elements as nodes, and regulatory relationships as edges. Transcription factors (TFs), a major class of protein regulator in gene expression (Thompson et al., 2015), are pivotal regulatory factors in these networks. Under different cellular conditions, TFs may undergo dramatic functional changes (or “network rewiring”) according to the gains and losses of their regulatory target genes. These rewiring events provide insight into differential cellular responses across conditions in the form of altered regulatory programs. Studies have revealed that network rewiring events and the altered regulatory programs they generate have strong phenotypic impacts (Bhardwaj et al., 2010, Assi et al., 2019).

However, quantification of network rewiring is challenging due to regulatory network’s condensed and complex nature (Gerstein et al., 2012). Genes from various functional modules, pathways, and molecular complexes can play varying roles depending on their local associations with other genes. As a result, gains or losses of some gene connections may impact network alterations in a functionally significant way, while others may not. This indicates that identifying the gene functional subgroups and estimating the network rewiring from the subgroup level should be more robust and informative, as compared investigating change of every individual gene. Thus, a low-dimensional representation of the regulatory network is required.

These low dimensional representations of functional subgroups underlying network data resemble semantic topics in documents. Based on this consideration, the low-dimensional representation can be constructed using topic modelling techniques including latent Dirichlet allocation (LDA). LDA was proposed by Pritchard et al. (Pritchard et al., 2000) for population genotype inference, and rediscovered by Blei et al. (Blei et al., 2003) with applications in natural language processing as a simple and efficient means to extract latent topics from high-dimensional data. It has been successfully implemented in several biological scenarios that require decomposition and dimensionality reduction of data (Pinoli et al., 2014, Wang et al., 2011). To apply LDA, we represent the targets of a TF under a specific condition (cell line or tissue) as a “document,” with the TFs’ target genes as “words” and latent functional subgroups as gene “topics” comprised of these words.

In this study, we propose a method called TopicNet that makes use of various features in the LDA model to measure the regulatory potential, perturbation tolerance, and intra-network dynamics of TFs in terms of their target gene topics. The training corpus includes all the regulatory networks inferred from 863 chromatin immunoprecipitation-sequencing (ChIP-seq) assays of the ENCODE dataset.

We first apply an LDA model to the data to characterize the gene “topics” in an unsupervised fashion. From the trained model we could infer the topic component as the distribution over words for each topic, and the topic weight as the distribution over the topics for a given document. The topic component can be further annotated for biological significance including their relevance to certain biological pathways and processes. The topic weights of each document can be used to quantify the rewiring between two cellular conditions, such as different cell types or time points after treatment. Lastly, we define a topic activity score by incorporating the cell-specific expression of target genes and topic composition and characterize specific topics with expression scores associated with patient survival of several cancer types.

In summary, our framework provides a straightforward quantitative representation of TF regulatory network with biological significance, which could be further applied to many downstream analyses.

## Results

### TopicNet framework

LDA model is used to decompose the high-dimensional regulatory network into a selected number of latent topics related to certain biological pathways (Fig 1a,b). Based on the results, we constructed the TopicNet framework including two parts: topic rewiring score and topic activity score (Fig 1c,d).

**Fig 1.**
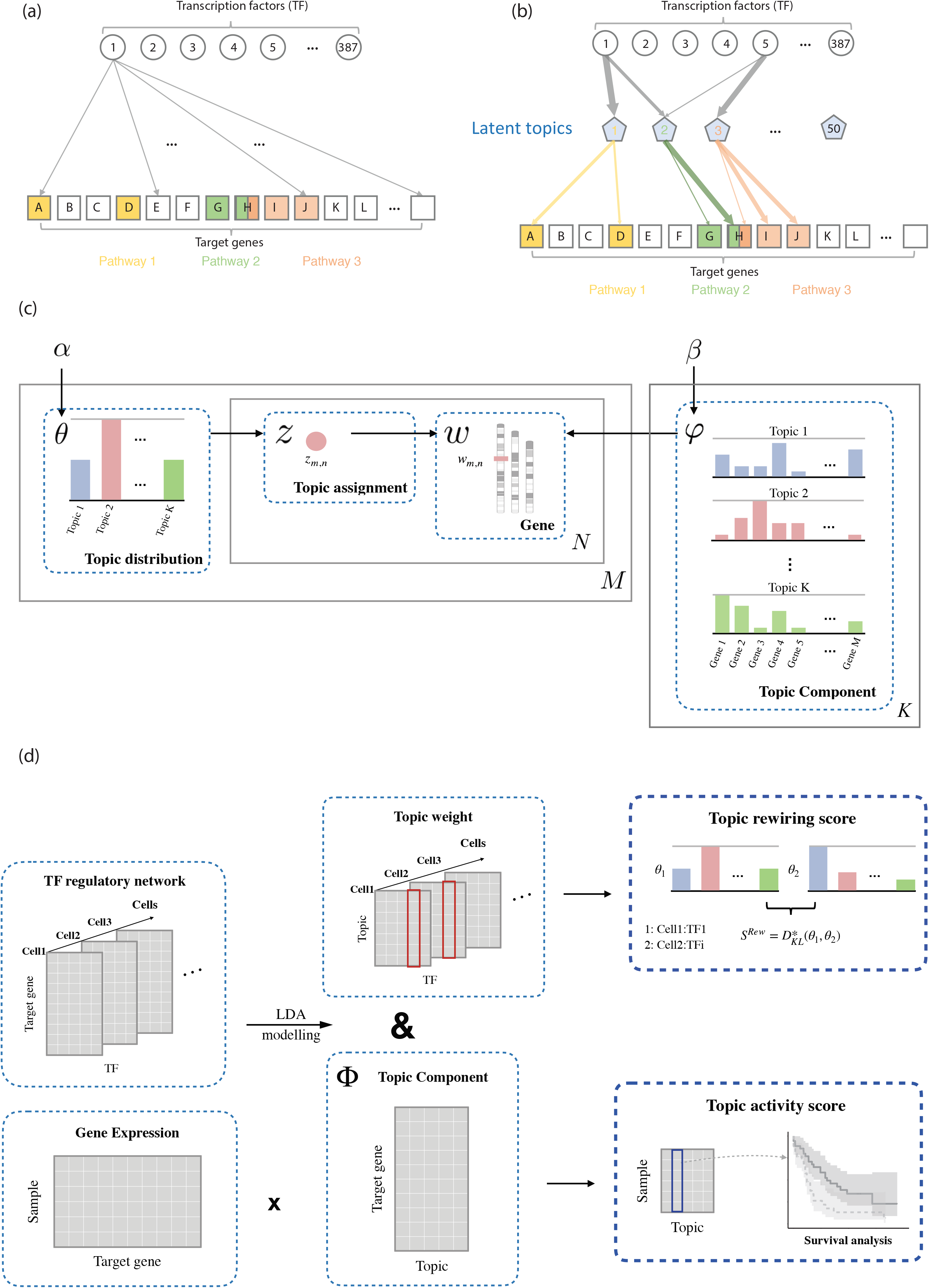
Overview of method and data. (a) An example of TF-gene regulatory network. (b) The high-dimensional regulatory network is decomposed into latent topics related to certain biological pathways. (c) Diagram explaining the meanings and biological relevance of the parameters in the LDA model. (d) General workflow of our analytical framework.

#### LDA model

In our study, we treated the regulatory targets of a TF under a specific cellular condition as a document, denoted as *W*_{*TF,cell*}_. The target genes act as “words” and constitute the general “vocabulary” of the corpus. The LDA then identified the functional topics from the genes in these documents (as described in Fig 1c). We used published metrics to choose the number of topics K; in particular, we use K=50 as the optimal number for our model (Fig s1; see “Methods” for more details).

Two important matrices can be inferred from the trained model, and their meanings are as follows:

1. The document-topic weight matrix Θ, which is cellular condition-dependent, represents the weight of topics for all documents. Each column θ_{*TF,cell*}_ is a vector of the distribution over topics for the corresponding document, and the element θ_{*TF,cell*},*k*_ represents the weight of topic *k* in document *W*_{*TF,cell*}_.
2. The topic-gene component matrix Φ, which is cell-independent, indicates the distribution of topic over the target genes. Each column *ϕ*_*k*_ is the component of the topics, i.e. the distribution of target genes to the topic or the contribution of genes to the topic. *ϕ*_*k,j*_ represents the contribution of gene *j* to topic *k*.

We further developed topic rewiring score and topic activity score based on these two matrices.

#### Topic Rewiring score

Raw network rewiring can be described as the differences between documents of the same TF in two cell types, *W*_{*TF,cell1*}_ and *W*_{*TF,cell2*}_. For comparison between two documents in terms of topics, we define network rewiring score as 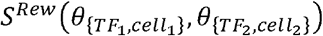 as a symmetrized Kullback-Leibler (KL) divergence between the topic distributions 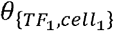 and 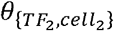.

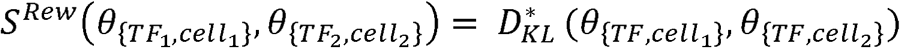

#### Topic activity score

Note that Θ gives an effective weighting to topics in a given condition. However, often the cell type-specific regulatory network is lacking, but gene expression is more available. In these cases, we can define an effective topic activity score. Given a sample *t* with gene expression vector *E*_*t*_, we compute the vector of expression score for all topics by multiplying It with the component matrix Φ, i.e. 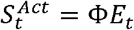.

### Validation of LDA model

We determined T=50 as an optimal number of topics using several metrics (Arun et al., 2010a, Cao et al., 2009, Griffiths and Steyvers, 2004). We then tested how similarities in the original data can be preserved compared with two other algorithms, non-negative matrix factorization (NMF) and K-means, using methods presented by Guo et al., 2017 (Guo and Gifford, 2017). Each method gives a 50-dimensional representation of the samples. For every pair of samples, we computed their correlation in terms of both the raw data (“raw” correlation) and the 50-dimensional representation (“reconstructed” correlation) from each method. Among the three algorithms, LDA could reconstruct the raw correlations better than the other two. (Fig 2a and Table 1). T-distributed stochastic neighbor embedding (T-SNE) of the 50-dimensional representation also demonstrates LDA’s ability to preserve similarities because samples about the same TF tend to form distinct clusters in the embedding space (Fig 2b).

**Fig 2.**
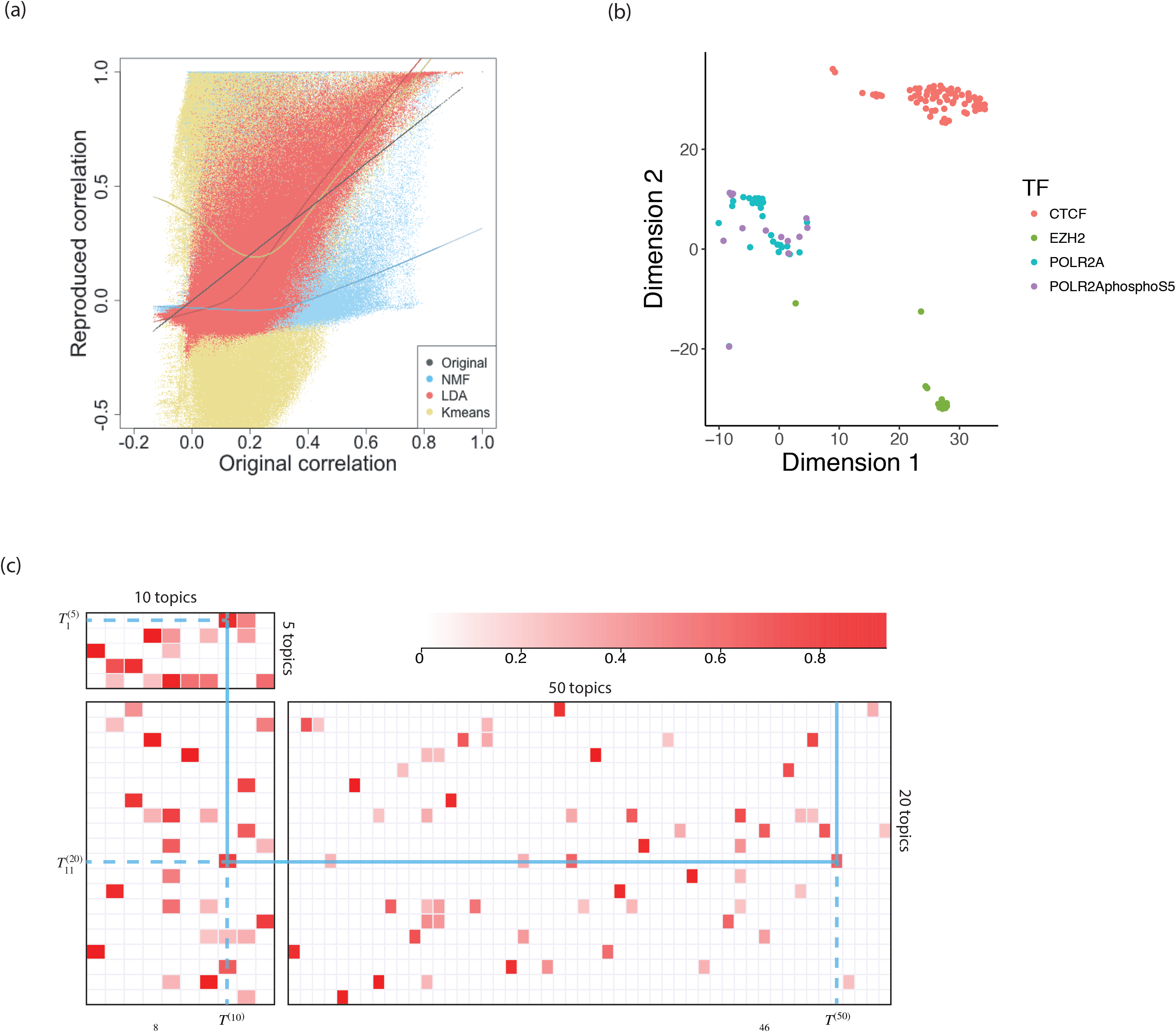
Tuning and performance evaluation of the LDA model. (a) Reproduced pairwise correlations after applying three dimensionality reduction methods plotted against original correlations. (b) T-SNE embedding of topic distributions (50 topic model) for data samples on CTCF, EZH2, POL2A and POL2AphosphoS5. (c) Correlation between topic components identified by LDA models with different topic numbers (5, 10, 20, 50).

**Table 1.** Linear regression of the reconstructed correlation against the raw correlation

| Method | Correlation | Linear regression slope | Linear regression $R^2$ |
| --- | --- | --- | --- |
| K means | 0.1047 | 0.3828 | 0.0110 |
| NMF | 0.4047 | 0.4365 | 0.1638 |
| LDA | 0.7886 | 1.3519 | 0.6219 |

We also investigated the connection of model that inferred by the different topic number *T*. We associated topics from models with the topic number *T* = 5, 10, 20,50 based on the correlation of their topic-gene component matrix and observed a natural hierarchical structure among these models (Fig 2c). This demonstrates that changing the topic number adjusts the “resolution” of the pattern recognition process as a model with more topics tends to detect more detailed patterns, while one with fewer topics can detect higher-level ones.

The topics can be associated with protein complex and functional modules. We performed hierarchical clustering on all TF documents in HeLa using their topic weights. Distinct clusters can be observed indicating co-activation and collaborative binding (Fig s2). Among them, nuclear transcription factor (NFY) subunits have been shown to co-localize with Fructooligosaccharides (FOS) extensively (Fleming et al., 2013), and FOS, NFYA and NFYB are clustered together. Similar grouping is also observed for CTCF and cohesin subunits SMC3 and RAD21, as the latter frequently co-bind with CTCF (Rubio et al., 2008, Parelho et al., 2008).

### Functional annotation of identified gene topics

We calculated the importance of each topic by measuring the Kullback-Leibler (KL) divergence between the topic weight of all documents against a background uniform distribution. We expect important topics should be more specific and only highly represented in some documents. The topics that are close to a uniform distribution have almost equal weights in most documents and will be less interesting. The rank of all 50 topics’ importance is shown in Fig 3a. Of particular interest are Topic 3 and 14: from the top genes of these two, we identified several functional groups such as transcription regulation, cell proliferation, metabolism and mitosis (Fig 3b).

**Fig 3.**
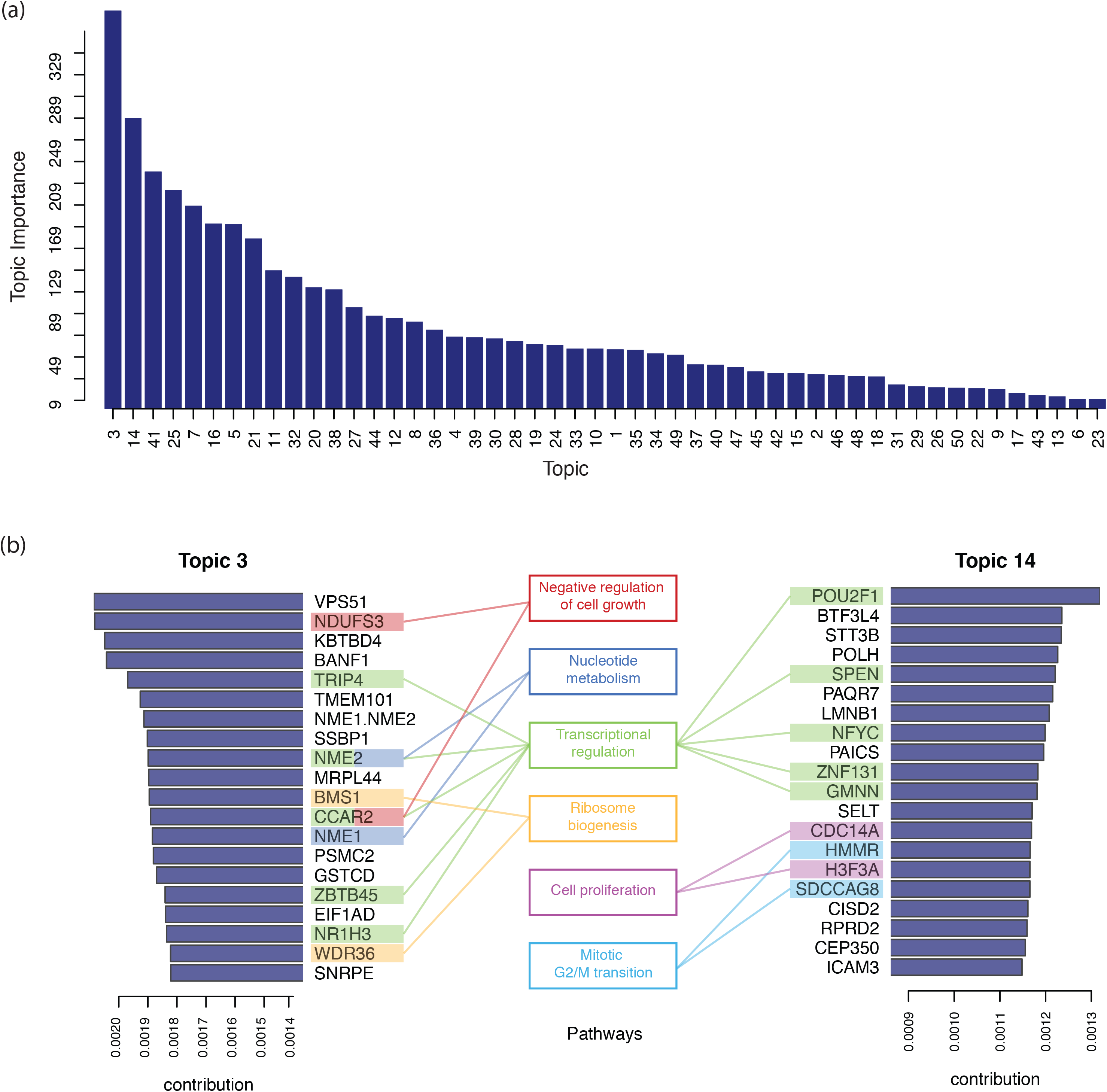
Annotation of the identified topics. (a) All 50 topics ranked by their importance measure (average KL divergence of topic distribution against uniform distribution across all samples). (b) Genes with highest contributions to the top ranked topics (Topic 3 and 14) along with their related functional roles.

To investigate the biological significance of each topic, we annotated their functions using gene set enrichment analysis (GSEA). For each topic, the probability distribution over target genes can be used directly as the statistics for GSEA. Using C2 and C5 gene sets from the Molecular Signatures Database (MsigDB) (Subramanian et al., 2005), Topic 3, the one with the highest importance, is enriched with gene sets related to breast cancer and glioblastoma tumors (Fig s3).

### Quantification of TF regulation rewiring using topic weights

For each TF, we calculated the topic rewiring score for every pair of cell type. The average rewiring score for a TF reflects its cell type specificity, as higher values correspond to greater difference between cell lines, i.e. higher specificity (Fig 4a, Fig s4). It can be observed that many TFs with higher cell specificity relate to biological processes displaying highly variable regulatory activity across conditions, such as pluripotency, cell cycle regulation, tumor suppression or tumorigenesis, including EP300 (Kim et al., 2013), BCL11A (Khaled et al., 2015, Dong et al., 2017), ZBTB33 (Pozner et al., 2016) and JUND (Caffarel et al., 2008, Millena et al., 2016). On the contrary, TFs with more constant roles such as NR2C2 (O’Geen et al., 2010) show very little difference between cell types. Interestingly, ZNF274 and SIX5, which were shown to relate to CTCF binding sites (Hong and Kim, 2017), also have low specificity similar to CTCF. Fig s5 lists the individual rewiring events with top values. Many of these events involve TFs with high cell type specificity, such as EP300, SUZ12, ZBTB33 and FOS.

**Fig 4.**
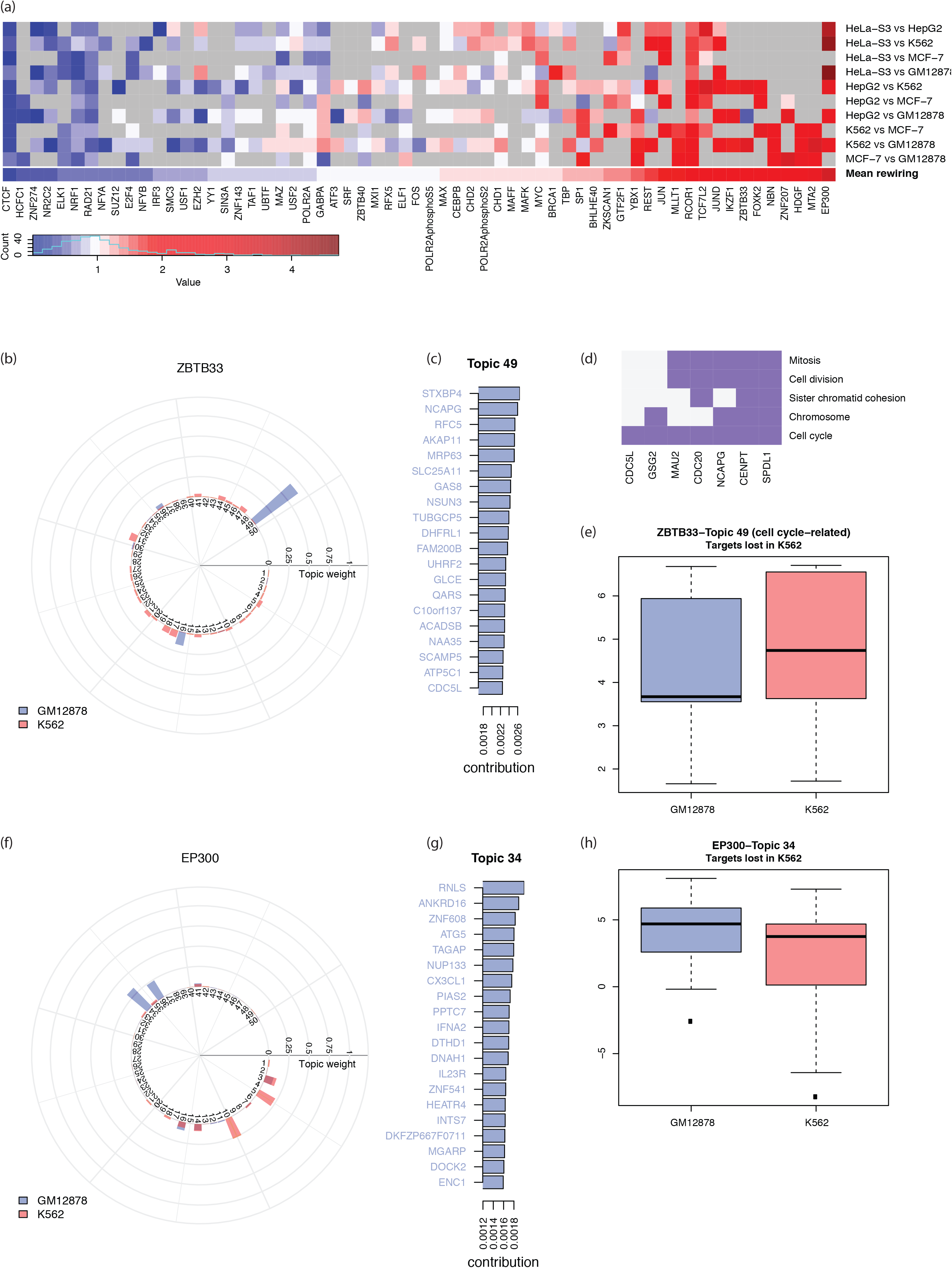
Quantified rewiring analysis using the identified topics. (a) Heatmap of rewiring of TFs in selected cell lines against GM12878 (grey grids correspond to unavailable data). (b) Topic weight for the rewiring event of ZBTB33 in GM12878 and K562. (c) Top weighted genes of the two topics with greatest difference in the rewiring event of ZBTB33 in GM12878 and K562. Colored gene names indicate true regulatory targets in the ChIP-seq experiment. (d) Functional clustering of top-rank genes of Topic 49 that are related to cell cycle and cell division. (e) Expression of the top-rank genes of Topic 49 which are lost in K562. (f) Topic weight for the rewiring event of EP300 in GM12878 and K562. (g) Top-rank genes of the two topics with greatest difference in the rewiring event of EP300 in GM12878 and K562. Colored gene names indicate true targets in the ChIP-seq experiment. (h) Expression of the top-rank genes of Topic 34 which are lost in K562.

We pinpointed two cell lines, GM12878 and K562, and studied specific rewiring events for several TFs. Among the 69 TFs shared in both cell lines, ZBTB33 and EP300 show the highest rewiring values. Specifically, Topic 49 and 16 show the greatest difference in the rewiring of ZBTB33 (Fig 4b), and Topic 34 and 10 in that of EP300 (Fig 4f). For these topics, the majority of their top-rank genes are true targets in the respective cell line (Fig 4c, g, Fig s6).

Given a specific TF, we defined “gain” target genes in K562 compared to GM12878 as those that are exclusively present in the former, and “loss” target genes as vice versa. In this scenario, we were particularly interested in the gain or loss genes among those with high contribution to the topic, i.e. the top-rank genes, that show major difference.

For ZBTB33, Topic 49 showed a very high weight in GM12878. We found the top-rank loss genes in K562 for Topic 49 is enriched in the gene set related to cell cycle and cell division function (Fig 4d). The loss of ZBTB33 regulation results in higher expression of its target genes in K562 corresponding to known de-acylation and transcriptional suppressive roles of ZBTB33 (Pozner et al., 2016). (Fig 4e). Another highly rewired TF, EP300 (a known transcriptional activator) regulates a wide range of genes from different functional groups. In concordance with EP300’s function, Topics 5 and 10 are deficient in K562 and the top-rank loss genes from these topics are significantly down-regulated in K562. For topics of EP300 that are highly represented in K562 (34, 36), an adverse trend is observed (Fig 4h, Fig s6). To summarize, topics showing major difference in the rewiring of these TFs are related to the TFs’ molecular functions. Comparatively, transcription factors with low rewiring score like CTCF and ZNF274 have almost identical topic distributions (Fig s7).

These results further demonstrate the potential of rewiring score derived from LDA as a quantitative measure of changes. The rewiring events with high scores could be associated with previously reported biological significance of corresponding TFs.

### Network rewiring shows dynamics topic changes of time-course study

Temporal changes of topic weights could be used to represent dynamic responses in the cellular regulatory system. To demonstrate this, we further applied our methods to the time series TRNs for estrogen receptor (ESR1) in MCF-7 cell lines at 2min, 5min, 5min, 10min, 40min and 160min after estradiol treatment (Guertin et al., 2014). Rewiring score between these time points shows transition of the topic distributions across time points. The first few minutes right after estradiol treatment have much dramatic topic changes and then it gradually go stable (Fig 5a).

**Fig 5.**
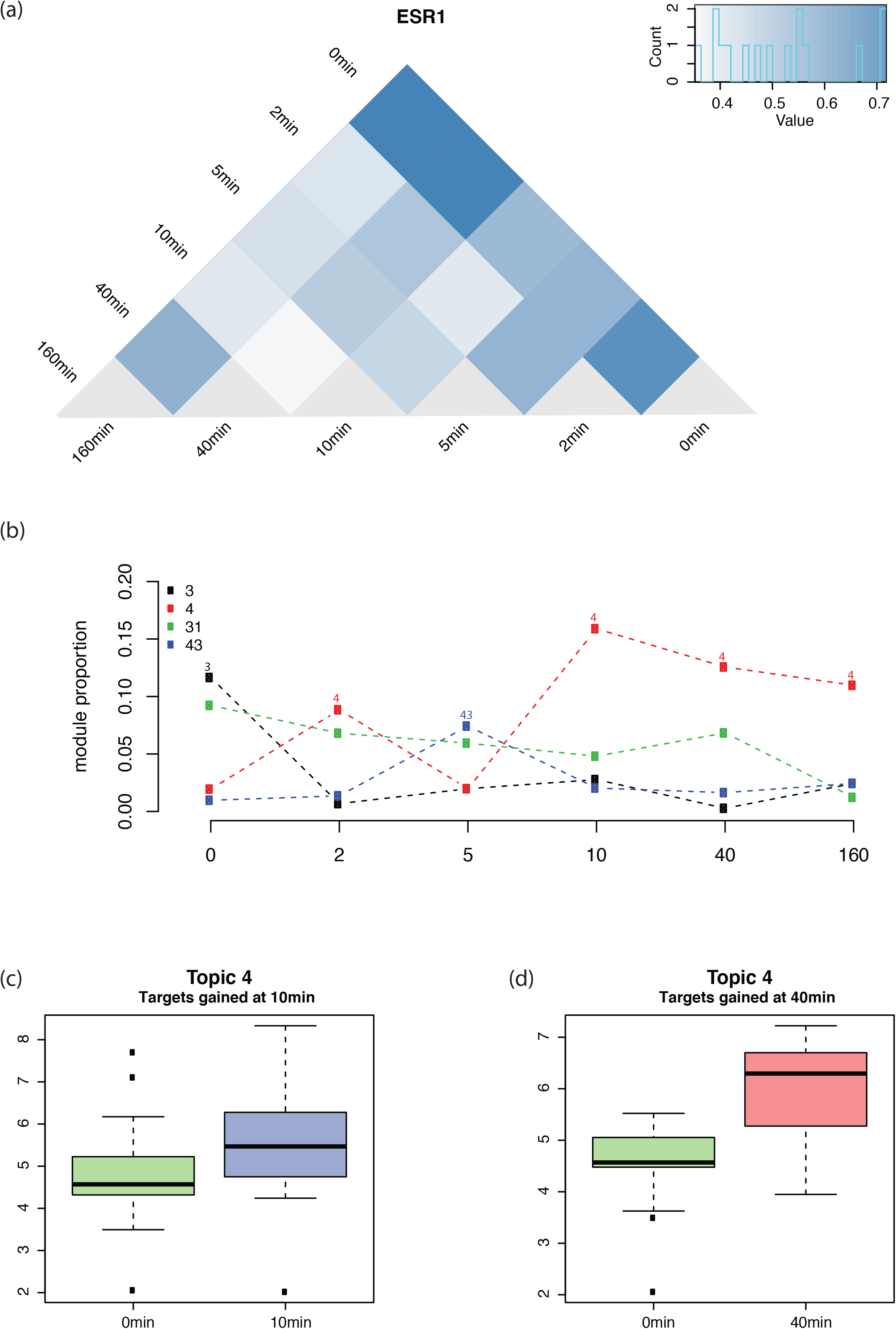
ESR1 regulation dynamics represented by topic distributions. (a) Pairwise KL divergence between the topic distributions of ESR1 in all time points. (b) Time course change of the topics with highest weights across the time points. (c)~(d) Expression of the genes gained in 10min (c) and 40min (d), as compared to 0min, which are among the top-rank genes of Topic 4.

The time course pattern of the topic membership demonstrates that Topic 3 has the highest weight prior to the treatment. Later, Topic 4 becomes the most prominent topic after short fluctuation, which experiences a sharp increase at 10 min time point and then undergoes a gradual decrease but still keep the dominant position until the end (160m) (Fig 5b). We then studied the roles of ESR1 target genes that are related to these highly represented topics. At 0min, true ESR1 target genes that are top-rank in Topic 3 include genes are most related to cell proliferation functions: ribosomal functions, protein folding, and mRNA splicing (Fig s8). At the 10min time point for Topic 4, top-rank target genes include genes related to signal transduction and apoptosis, with some of them interacting directly with EP300, which is consistent with the findings that the redistribution of EP300 target after the treatment of estradiol (Guertin et al., 2014) (Fig s8).

The treatment of estradiol turned the MCF-7 cell line’s topic from Topic 3 to 4, which indicates the top weighted genes in topic 4, especially for the gained genes, may play a crucial role in the treatment. We compared the nascent gene expression of 10-min and 40-min with 0-min for gained top-rank gene in topic 4. These gained top-rank genes show very significant (p-value < 10^−5^) up-regulation in 10-min and 40min (Fig 5c, d).

### Topic activity score and its relationship to tumor survival

Topic activity score incorporates cell type-independent topic component with cell type-specific gene expression and can be associated with clinical significance. We used patient samples of 3 cancer types with clinical information from the TCGA data portal breast cancer (BRCA), acute myeloid leukemia (LAML) and liver hepatocellular carcinoma (LIHC). Topic activity scores were evaluated for each cancer type and used for survival analysis. We found several topics whose expression score is associated with patient survival (Fig 6). For each cancer type, we characterized the biological relevance of the most predictive topic:

1. Expression score of Topic 10 is predictive for the survival of BRCA patients (Fig 6a). Correspondingly, Topic 10 is highly represented in the document of (Zhang et al., 2017), and its component is enriched with CtIP associated gene set (Fig s9a). GATA3 and CtIP are known to interact with each other and functionally correlate with breast cancer: GATA3 can regulate BRCA1 (Zhang et al., 2017) and CtIP forms a repressor complex with BRCA1 whose removal accelerates tumor growth (Furuta et al., 2006) (Fig s10a).
2. Expression score of Topic 26 is predictive of LAML patient’s survival. Topic 26, which is highly represented in the document {NR2C2, K562} (Fig 6b, Fig s9b), its components is also enriched with genes up-regulated in response to activation of the cAMP signaling pathway (van Staveren et al., 2006) (Fig s10b). NR2C2 can be induced by cAMP (Liu et al., 2009) and are found to be significantly active expressed in almost all the cancers (Falco et al., 2016).
3. Expression score of Topic 35 predict survival outcome of LIHC patients with high accuracy. Topic 35 is highly represented in the documents {ATF3, HepG2} and {JUN, HepG2} (Fig 6c, Fig s9c), and its component is enriched with gene set that are up-regulated in response to over-expression of proto-oncogene MYC (Bild et al., 2006) (Fig s10c). Among these factors, ATF3 is a cAMP-responsive element and acts as a tumor suppressor in LIHC (Chen et al., 2018). JUN is a known oncogene and promotes liver cancer (Maeda and Karin, 2003). MYC is also a highly expressed oncogene and correlates with high proliferative activity (Zheng et al., 2017).

**Fig 6.**
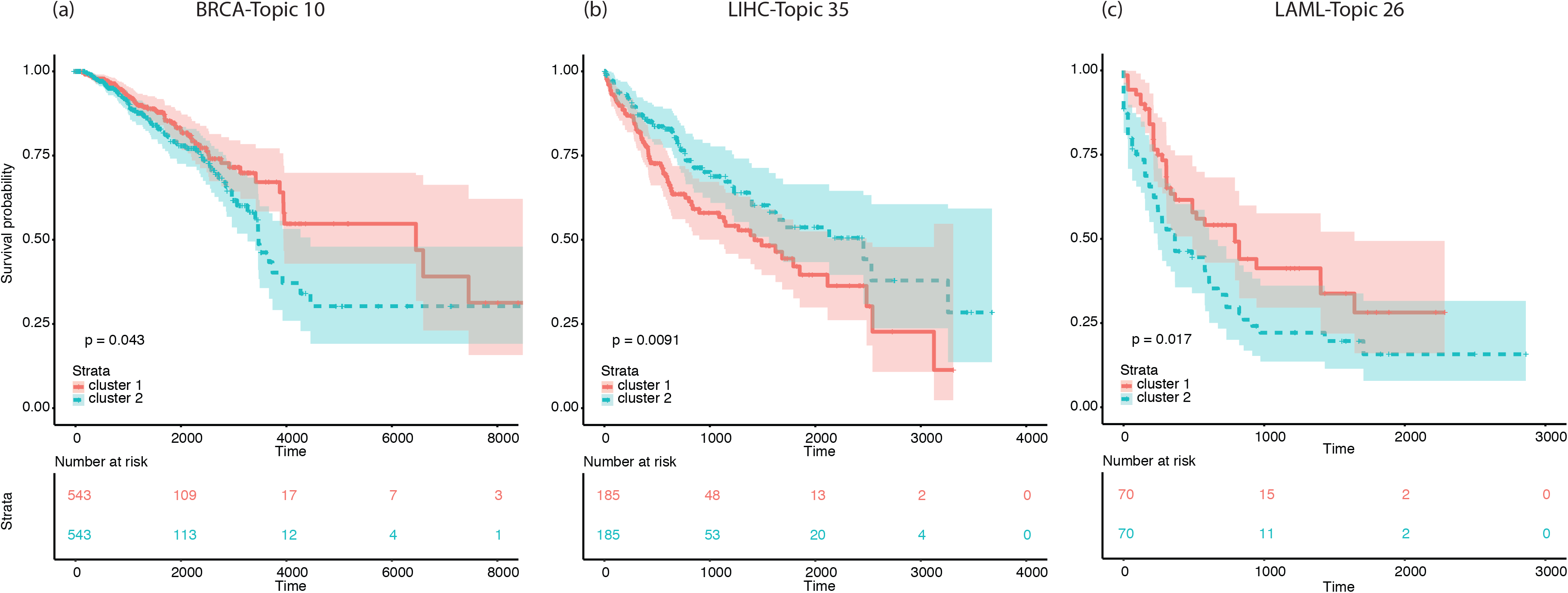
Topic activity level related to cancer survival. (a)~(c) Kaplan-Meier survival curve of thee cancer types using their related topic activity levels: BRCA with topic 10 (a), LIHC with Topic 35 (b) and LAML with Topic 25 (c).

In summary, we have found associations between the survival-related topics and their biological significance via the activity and function of the TFs that regulate these topics. These results further validate the biological relevance of the identified topics, indicating their potential as prognostic markers and sources for biomarker discovery.

## Discussion

Rewiring analysis of the regulatory network could provide critical information about the alteration of molecular programs across conditions. Several attempts have been made to derive an effective procedure for identification of network rewiring. In this study, we successfully developed a TopicNet framework. Our framework extracted a low-dimensional representation of network in the form of functional topics, defined network rewiring score and topic-weighted expression score, and then we demonstrated the application of the framework. The network rewiring score can aid in the identification of the functional rewiring of TFs between different cellular conditions. The topic-weighted expression score can be applied to sample-specific cohort data for the prediction of patient survival.

As a verification of our framework, we also investigated the biological meaning of the identified topics. We interpreted the learned topics by utilizing the two important matrices inferred from the model: document-topic weight and topic-gene component matrices. The former demonstrates the activity of the topic as a distinctive functional module, and the latter indicates possible biological functions or pathways that the topic represents. Rewiring analysis using topic weights is both efficient and highly interpretable with gene topics serving as bridges between TFs and genes.

Our framework facilitates comparison between regulatory networks under different conditions from multiple sources. Thus, the analysis can be extended to various studies where network changes are of major interest. For example, time-course network changes, such as those after treatment or during the cell cycle, could help pinpointing TFs, genes and pathways that play critical roles in these processes. Having demonstrated the potential application of our method in time course data, we expect our method to offer valuable insights into network dynamics studies in the future.

LDA has been shown to be advantageous over some other common dimensionality reduction techniques in terms of performance and interpretability (Stevens et al., 2012, Liu et al., 2011). Furthermore, several extensions on the LDA model could be introduced for future studies. In our framework, we treated all TF-cell line pairs as independent regardless of possible relationships between documents. Additionally, the low-dimensional gene topic defined by TopicNet is a simplified representation, which does not take into account the complex and hierarchical gene-gene interactions. We are aware that recent advances in topic modelling methods has enabled modelling of more complex dependencies and structures (Blei and Lafferty, 2006, Momeni et al., 2018, Zhou et al., 2017). Though LDA could capture some of these dependencies in an unsupervised fashion, we expect incorporation of such information would help identify even more meaningful patterns from the regulatory network.

## Supporting information

SuppFigures

## Author contributions

SL: conceptualization, methodology, formal analysis, visualization, writing-original draft; TL: methodology, formal analysis, visualization, writing-original draft; XK: methodology, formal analysis, visualization, JZ, JL and DL: resources, validation;MG: supervision

## Methods

### Data preprocessing and construction of the regulatory network

We used 863 ChIP-Seq experimental results for 387 TFs from the ENCODE portal for model training due to their higher quality control and consensus peak calling. In addition, we included ChIP-Atlas data collections with more than 6,000 ChIP-Seq experimental results to test the model. The number of target genes included in this dataset ranges from hundreds to thousands (Fig s11), and the TFs with the greatest availability among different cell lines include CTCF, EP300, MYC and REST (Fig s12).

From each ChIP-seq experiment, the regulatory target genes of specific TFs are defined as those with ChIP-seq peaks in proximal regions (+/− 2500bp) of their transcription start site. The cell type-specific TRN is then defined based on these results.

### TopicNet - Topic modelling

Each regulatory network for a TF in a specific cell line is regarded as an independent input document. We treat target genes that exist in these documents as “words”, which collectively constitutes the “vocabulary” of the model. Based on existence of all genes as a regulatory target of the TF in the given condition, a document-gene matrix is then constructed. This matrix is used as the input for the LDA model.

Let *M*, *K*, *V* be the number of documents, the number of topics and the vocabulary size, respectively. In this scenario, each document is modelled as a mixture of topics, and each topic is a probabilistic distribution over genes. Each document *t* is represented as a *N*_*i*_-dimensional vector *W*_*i*_, where *N*_*i*_ is the number of genes in the document, and each element takes the value 1…*V*. The probability of observing a gene *w*_*ij*_ in a document *W*_*i*_ is determined by the mixture of topic components within the document and the probabilistic distribution of those topics. The existence of a word in a document is modelled as follows (Fig 1a):

Given two priors (*α* as the prior for document-topic distribution, and *β* as the prior topic-gene distribution) we can sample

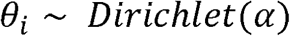

as the probability of all topics appearing in a document *i*, which constitutes the *M* × *K* matrix for document-topic distribution; and

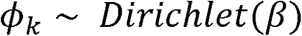

as the probability of all genes appearing in a topic *k*, which constitutes the *K* × *V* matrix for topic-gene distribution.

We can then sample latent topic assignment of each word *j* in document *i* as

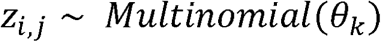

which is the topic that generates this gene. Each *z*_*i,j*_ can take the value 1 … *K*.

Given the membership of latent topics, the existence of genes in a document can be drawn as

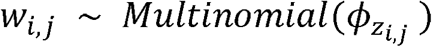

which constitutes the observed document-word matrix, where 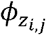 is the topic-word distribution for the sampled topic *z*_*i,j*_.

### Model inference

Let *W* and *Z* be the collection of all aforementioned *w*_*i,j*_’s and *z*_*i,j*_’s indexed by document and gene position pair (*i, j*). The model parameters can be estimated using a collapsed Gibbs sampler (Heinrich, 2005) on a Markov chain of {*W, Z*} where *W* is the observation and *Z* is the hidden variable. The joint distribution of the LDA model 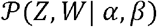 can be obtained by integrating out *ϕ*, and *θ*:

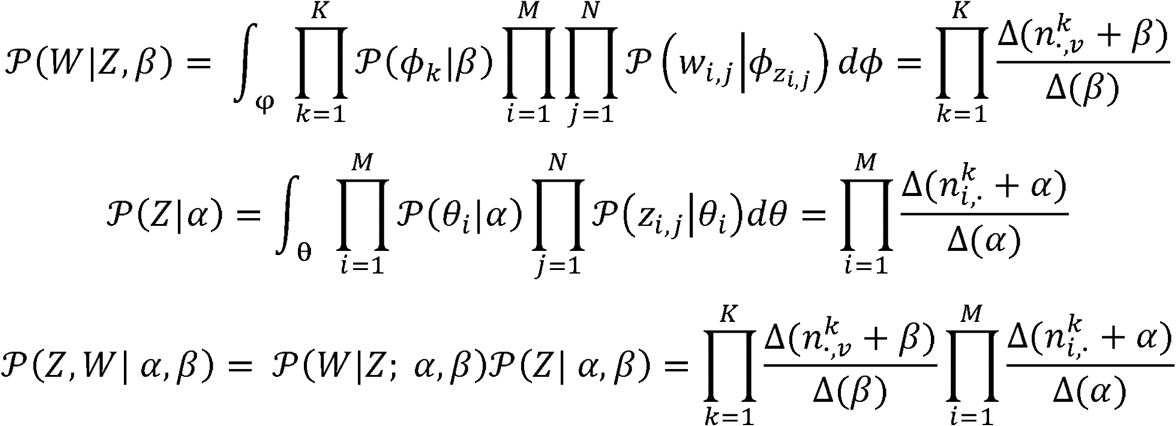

where 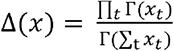 for a *t*-dimensional vector 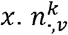 is the number of times gene *v* is assigned to topic *k* and 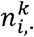 is the number of times genes in document *i* are assigned to topic *k*. These sums can all be calculated from the value of {*W, Z*}.

The conditional probability of *z*_*i,j*_, the topic assignment of the *j*-th gene in the *i*-th document, can be inferred as:

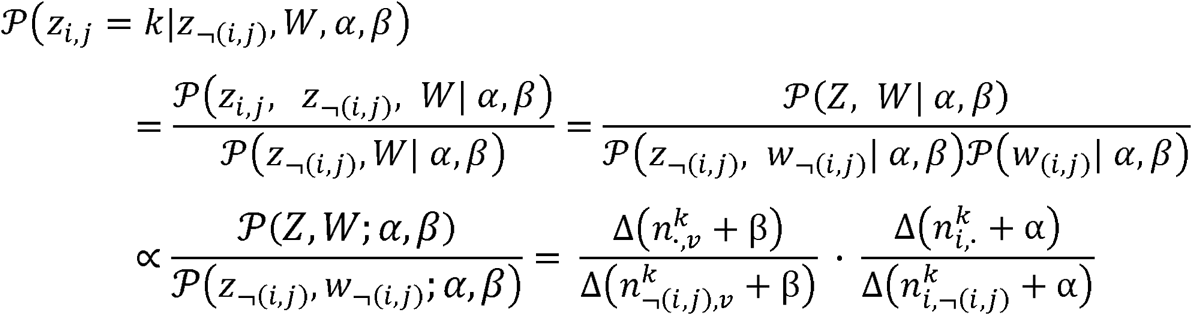

where 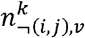 and 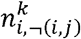 indicate the same type of count as mentioned above excluding (*i, j*).

This gives the sampling distribution for *Z* given *W*.

Using the above formulae, Gibbs sampling can then be performed on the Markov chain {*W, Z*} to obtain the estimation of *φ*, and *θ*. By definition, we have the probability distribution of {*W, Z*} conditioned on *φ* and *θ*. Using Bayes rule, the expectation of *φ* and *θ* can be inferred from {*W, Z*}:

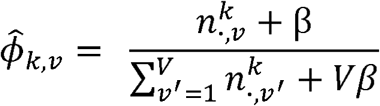

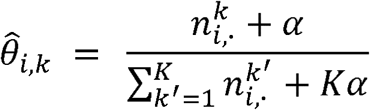

For a stable and robust topic-gene component matrix, we averaged the results of 100 runs. As the topics learned by the LDA model for each run were represented in randomized orders, topics across different samplings were first mapped against each other based on the correlations of their components. For any pair of outputs, each topic from the first run is assigned with the same ID as the topic it most strongly correlates with in the other run. We then produced the ensemble model by taking the median of the probability distribution over the components for all topics that are mapped to the same ID.

After we obtained the model, we can apply it to unseen documents with the same vocabulary and determine their posterior distribution over topics given the generative processes above.

### Selection of topic numbers

The number of topics *T* used was selected using multiple criteria: the posterior likelihood of the data given the LDA model of different choice of number (Griffiths and Steyvers, 2004) (blue); KL divergence of the document-gene matrix (Arun et al., 2010b) (red); and the average cosine distance r within topics (Cao et al., 2009) (green). An optimal *T* should result in higher likelihood and lower distances. All three metrics reached optimal performance at around 50 topics (Fig 2a). Based on these results, we used 50 as the number of topics for downstream analysis.

### Reconstructed correlations

We performed LDA, NMF and K-means with 50 topics on each sample to obtain one 50-dimensional embedding vector of each sample for all three models. We represent the raw data for document *i* as a vector *v*_*i*_ = [*v*_*i,1*_, *v*_*i,2*_, …, *v*_*i,N*_] with binary values where *N* is the number of all genes in the vocabulary. The embedding procedure for each method is as follows:

For LDA, we obtain the document-topic weight matrix Θ as described above. For each document *i*, the embedding vector is the weight of the 50 topics, which is the *i*-th column of matrix Θ:

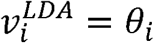

NMF decomposes the input matrix into two non-negative matrices: the feature matrix *W*, and the coefficient matrix *H*. For a document *i*, we use its weights as the embedding vector, which is the *i*-th column of matrix W:

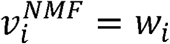

K-means identifies k=50 clusters from the dataset, and each cluster is represented as its cluster centroid 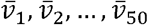, which is the average of all samples assigned to the respective cluster. Each document *i* is represented as the Euclidean distances between the raw vector and the 50 cluster centroids:

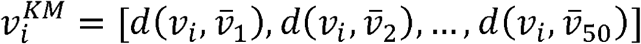

For document pair in the dataset, we could calculate their Pearson correlation using the raw vectors and the 50-dimensional embedded vectors of the three methods. To evaluate whether the embedding retains correlations in the original data, the “reconstructed” correlation calculated from the embedding vectors was then plotted against the “original” correlation using the raw vectors. Linear regression is also performed between the reconstructed and original correlation for the three methods respectively.

### Evaluation of topic importance

The importance of the ensemble topics is evaluated by calculating the KL divergence between the topic weights of all the documents and a background uniform distribution. Given the document-topic weight vector for all documents *θ* = [*θ*_1_, *θ*_2_, …, *θ*_*N*_](where N is the number of documents), the noise distribution is defined as:

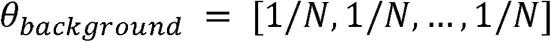

The distance between the ensemble distribution and the null distribution is defined as the KL divergence:

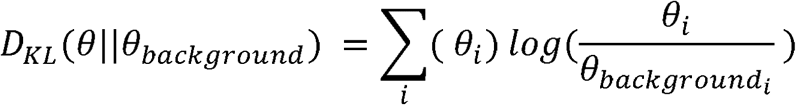

### Gene set enrichment analysis

We use topic component as the statistic for gene set enrichment analysis. Gene sets in the C2, C5 and C6 categories from MSigDB (Subramanian et al., 2005) were used in the analysis. Gene set enrichment analysis is performed with R package *fgsea*. Affinity between gene sets is defined by their overlapping of gene sets.

### Topic rewiring score - Quantification of network rewiring

For a pair of documents in a rewiring event of a given TF 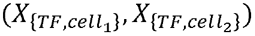, we can calculate the KL divergence between their topic weight vectors 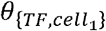 and 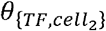, here using the later as reference:

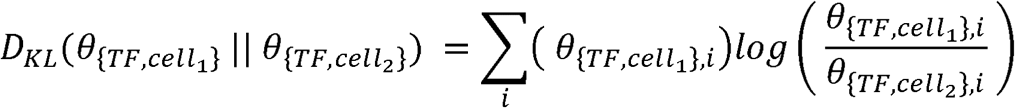

Since KL divergence is asymmetric depending on which distribution is used as reference, we consider the symmetrized KL divergence of the two directions to be a better metric for rewiring, which is:

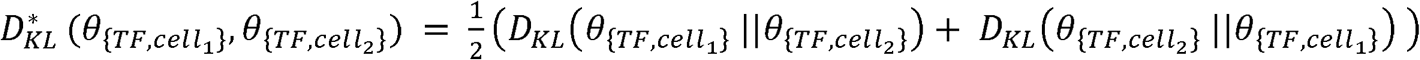

### Topic activity score

Gene expression data for BRCA, LAML, LIHC and GBM patient samples from TCGA are obtained from GDC data portal (https://portal.gdc.cancer.gov/). The gene expression levels of every sample in each cancer type is formulated into an expression matrix *E* where rows represent genes and columns represent samples. For each cancer type, the expression data is first quantile normalized. Only genes that appear in the topic-gene matrix are retained. We then obtain the expression matrix *E* by first ranking the expression of the genes for each sample and then transform the value for gene *i* in the matrix of column *j* (corresponding to patient sample *j*) to *E*_*i,j*_ = 1/*rank*_*i,j*_ where *rank*_*i,j*_ is the rank of gene *i* in sample *j*.

For a sample *t* with expression vector *E*_*t*_, the topic activity score is calculated as i.e. 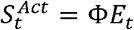, where each element in the vector is the expression score of the corresponding target.

#### Definition of gain or loss genes

Given two TF regulatory networks under two conditions, we arbitrarily assign one as an “altered condition” and the other as the reference condition. We define “gain” genes as those that are only regulated by the TF in the altered condition but not the reference condition, (formally called “gain genes in the altered condition”), and the loss genes vice versa (called “loss genes in the altered condition”). For annotation of selected topics, we are particularly interested in the gain or loss genes that are among the top-rank genes of the topic, i.e. those have high values in the topic component. We take the intersection between these two sets and name the resulting set of genes as “top-rank gain/loss genes in the altered condition for topic k”.

#### PPI network analysis of topic-related target genes

The genes in the corpus are first sorted according to their contributions to the topic. Among the top 500 genes, those that are directly regulated by the given TF (i.e. bound by the TF in the corresponding ChIP-Seq experiment) are selected and provided to STRING (Franceschini et al., 2013). The resulting interaction graph contains the selected genes along with their first-layer neighbours.

#### Survival analysis

Clinical information for each patient regarding vital status, days to last follow-up, and days to death are downloaded from GDC data portal (https://portal.gdc.cancer.gov/). Records with missing information are discarded. Patients that are still alive in the record are right censored. The values of all 50 topics are used as variables to perform Cox proportional hazards (coxph) regression and implemented with the coxph function from R package *survival*, with days to death (or days to last follow-up for censored living patients) as the response.

We then select the topics, S^Exp^ score of those have p-value < 0.05 in the coxph analysis, for further analysis. For each of these candidate topics, the patients are then separated into two groups by the median value of S^Exp^. The survival curve is then estimated using the Kaplan-Meier estimator. The topics that achieves lowest p-value is selected and shown.

## Supplementary Figure

**Fig s1** Selection of topic number using three different metrices.

**Fig s2** Hierarchical clustering of TF regulatory roles in HeLa-S3 using pairwise KL divergence as distance measure.

**Fig s3** Correlation of the highest represented gene sets in the component of Topic 3. Numbers indicate shared genes among the gene sets.

**Fig s4** All TFs ranked by their average pairwise rewiring values across cell types.

**Fig s5** Table of rewiring events with highest rewiring values.

**Fig s6** Topic component and expression of the top weighted genes that show highest rewiring in EP300 regulation between GM12878 and K562 after Topic 34: Topic 10 (a), 36 (b) and 5 (c).

**Fig s7** Topic weight of regulatory targets for the two lowest rewiring events: CTCF (left) and NR2C2 (right) in GM12878 and K562.

**Fig s8** Roles in protein-protein interaction networks of the top-rank target genes of Topic 3 at 0min (a) and Topic 4 at 10min (b). Deep blue nodes are the selected target genes and light blue nodes are their first-layer neighbors in the network.

**Fig s9** Gene set enrichment of oncogenic signatures (C6) gene sets in topics related to cancer patient survival (topic 10, 26 and 35).

**Fig s10** TFs in tumor cell lines that are highly enriched with survival-related topics: Topic 10 in MCF-7 (a), Topic 26 in K562 (b) and Topic 35 in HepG2 (c).

**Fig s11** Statistics of identified target genes from ChIP-Seq datasets used in the study.

**Fig s12** Log frequency of transcription factors with highest occurrence in ChIP-ATLAS and ENCODE datasets.

