## Supplementary material for "TopicNet: a framework for measuring transcriptional regulatory network change": SuppFigures

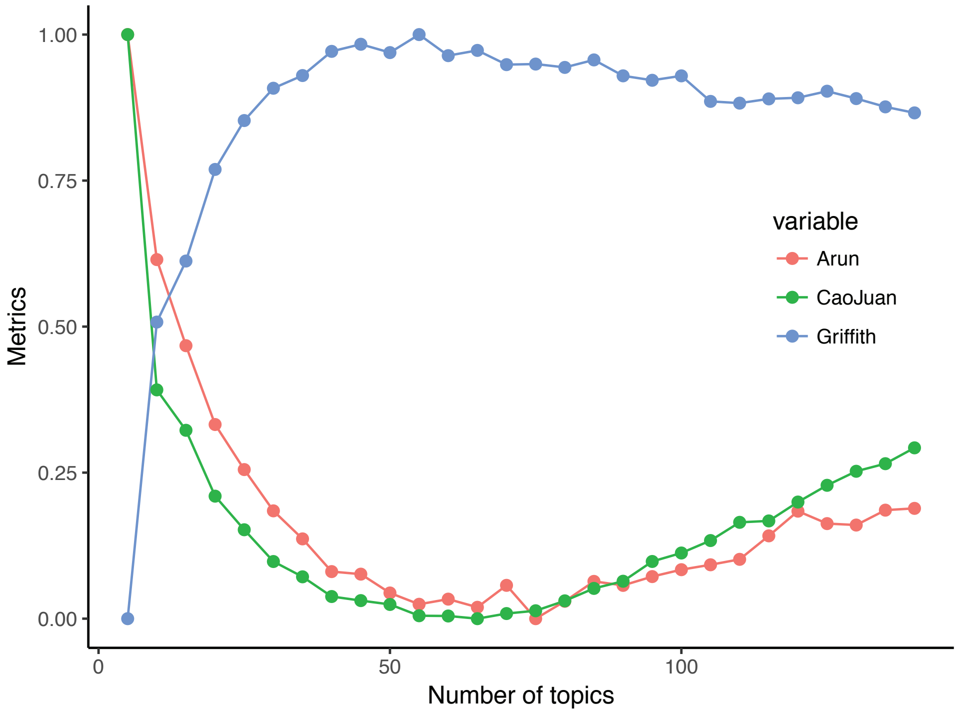

Metrics
1.00
0.75
0.50
0.25
0.00
0
50
100
Number of topics
variable
Arun
CaoJuan
Griffith

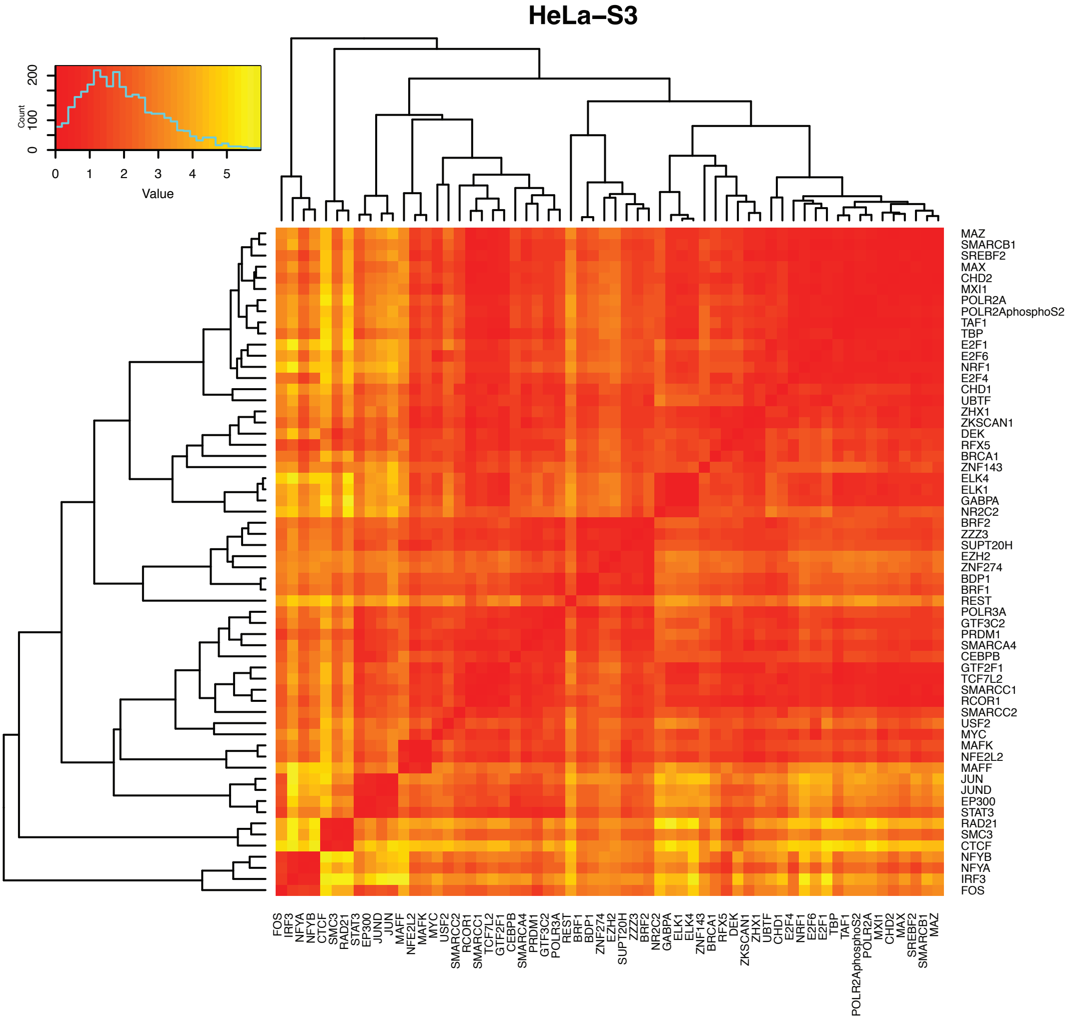
HeLa–S3
Count
0
100
200
0
1
2
3
4
5
Value
MAZ
SMARCB1
SREBF2
MAX
CHD2
MXI1
POLR2A
POLR2AphosphoS2
TAF1
TBP
E2F1
E2F6
NRF1
E2F4
CHD1
UBTF
ZHX1
ZKSCAN1
DEK
RFX5
BRCA1
ZNF143
ELK4
ELK1
GABPA
NR2C2
BRF2
ZZZ3
SUPT20H
EZH2
ZNF274
BDP1
BRF1
REST
POLR3A
GTF3C2
PRDM1
SMARCA4
CEBPB
GTF2F1
TCF7L2
SMARCC1
RCOR1
SMARCC2
USF2
MYC
MAFK
NFE2L2
MAFF
JUN
JUND
EP300
STAT3
RAD21
SMC3
CTCF
NFYB
NFYA
IRF3
FOS
FOS
IRF3
NFYA
NFYB
CTCF
SMC3
RAD21
STAT3
EP300
JUND
JUN
MAFF
NFE2L2
MAFK
MYC
USF2
SMARCC2
RCOR1
SMARCC1
TCF7L2
GTF2F1
CEBPB
SMARCA4
PRDM1
GTF3C2
POLR3A
REST
BRF1
BDP1
ZNF274
EZH2
SUPT20H
ZZZ3
BRF2
NR2C2
GABPA
ELK1
ELK4
ZNF143
BRCA1
RFX5
DEK
ZKSCAN1
ZHX1
UBTF
CHD1
E2F4
NRF1
E2F6
E2F1
TBP
TAF1
POLR2AphosphoS2
POLR2A
MXI1
CHD2
MAX
SREBF2
SMARCB1
MAZ

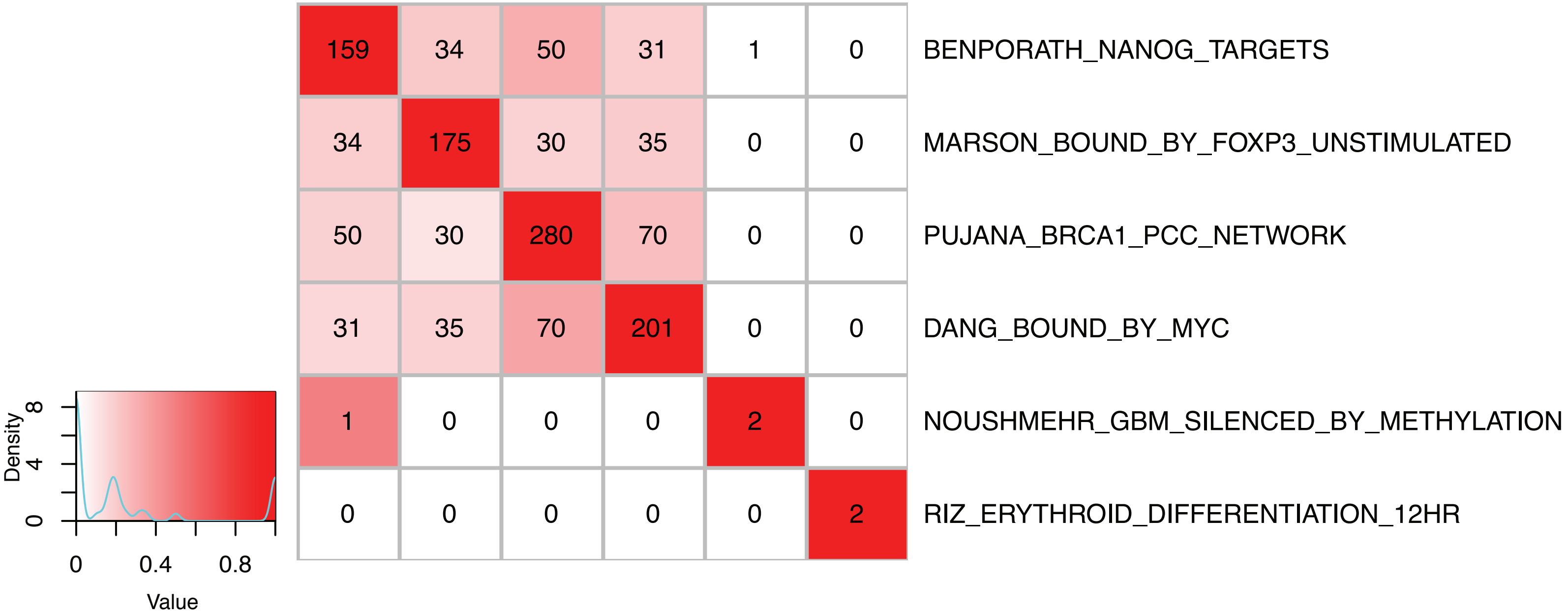

159 34 50 31 1 0 BENPORATH_NANOG_TARGETS
34 175 30 35 0 0 MARSON_BOUND_BY_FOXP3_UNSTIMULATED
50 30 280 70 0 0 PUJANA_BRCA1_PCC_NETWORK
31 35 70 201 0 0 DANG_BOUND_BY_MYC
1 0 0 0 2 0 NOUSHMEHR_GBM_SILENCED_BY_METHYLATION
0 0 0 0 0 2 RIZ_ERYTHROID_DIFFERENTIATION_12HR
Density
0
4
8
0
0.4
0.8
Value

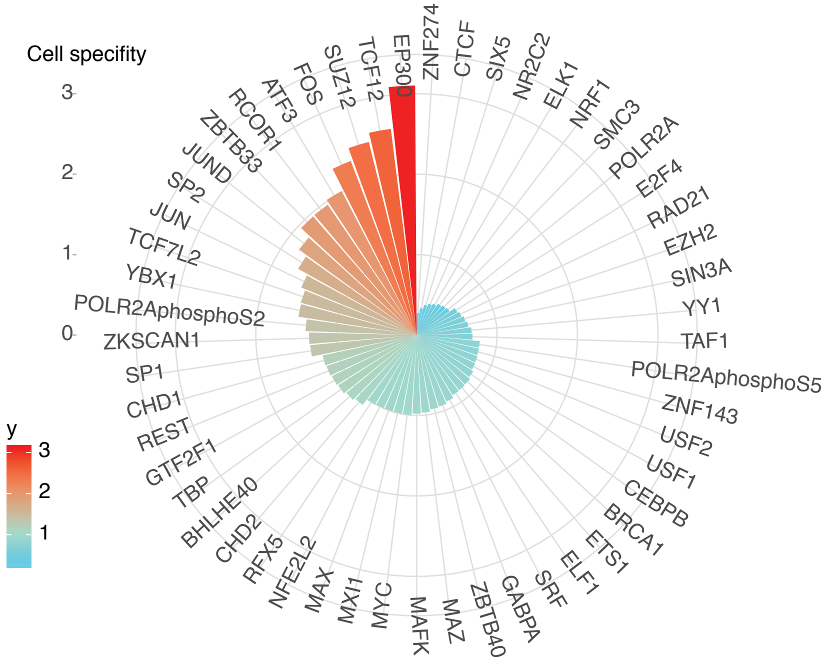
Cell specifity
3
2
1
0
y
3
2
1
EP300
ZNF274
CTCF
SIX5
NR2C2
ELK1
NRF1
SMC3
POLR2A
E2F4
RAD21
EZH2
SIN3A
YY1
TAF1
POLR2AphosphoS5
ZNF143
USF2
USF1
CEBPB
BRCA1
ETS1
ELF1
SRF
GABPA
ZBTB40
MAZ
MAFK
MYC
MXI1
MAX
NFE2L2
RFX5
CHD2
BHLHE40
TBP
GTF2F1
REST
CHD1
SP1
ZKSCAN1
POLR2AphosphoS2
YBX1
TCF7L2
JUN
SP2
JUND
ZBTB33
RCOR1
ATF3
FOS
SUZ12
TCF12

| TF | Cell–1 | Cell–2 | Rewire |
|---|---|---|---|
| FOS | GM12878 | endothelial–cell–of–umbilical–vein | 5.2951 |
| SUZ12 | HepG2 | NT2–D1 | 5.1265 |
| SUZ12 | H1–hESC | HepG2 | 4.7336 |
| EP300 | GM12878 | HeLa–S3 | 4.7269 |
| EP300 | HeLa–S3 | K562 | 4.4744 |
| ZBTB33 | GM12878 | HEK293 | 4.3908 |
| SUZ12 | K562 | NT2–D1 | 4.2648 |
| ZBTB33 | HEK293 | HepG2 | 4.161 |
| FOS | K562 | endothelial–cell–of–umbilical–vein | 4.1389 |
| EP300 | H1–hESC | HeLa–S3 | 4.0359 |
| EP300 | HeLa–S3 | HepG2 | 4.0125 |
| SUZ12 | H1–hESC | K562 | 3.942 |
| ZBTB33 | HEK293 | liver | 3.6158 |
| FOS | GM12878 | MCF–10A | 3.591 |
| TCF7L2 | HepG2 | K562 | 3.4521 |
| ATF3 | H1–hESC | liver | 3.3157 |
| TCF7L2 | HeLa–S3 | HepG2 | 3.3152 |
| ZBTB33 | A549 | HEK293 | 3.3096 |
| EP300 | A549 | GM12878 | 3.2969 |
| ATF3 | A549 | HepG2 | 3.269 |

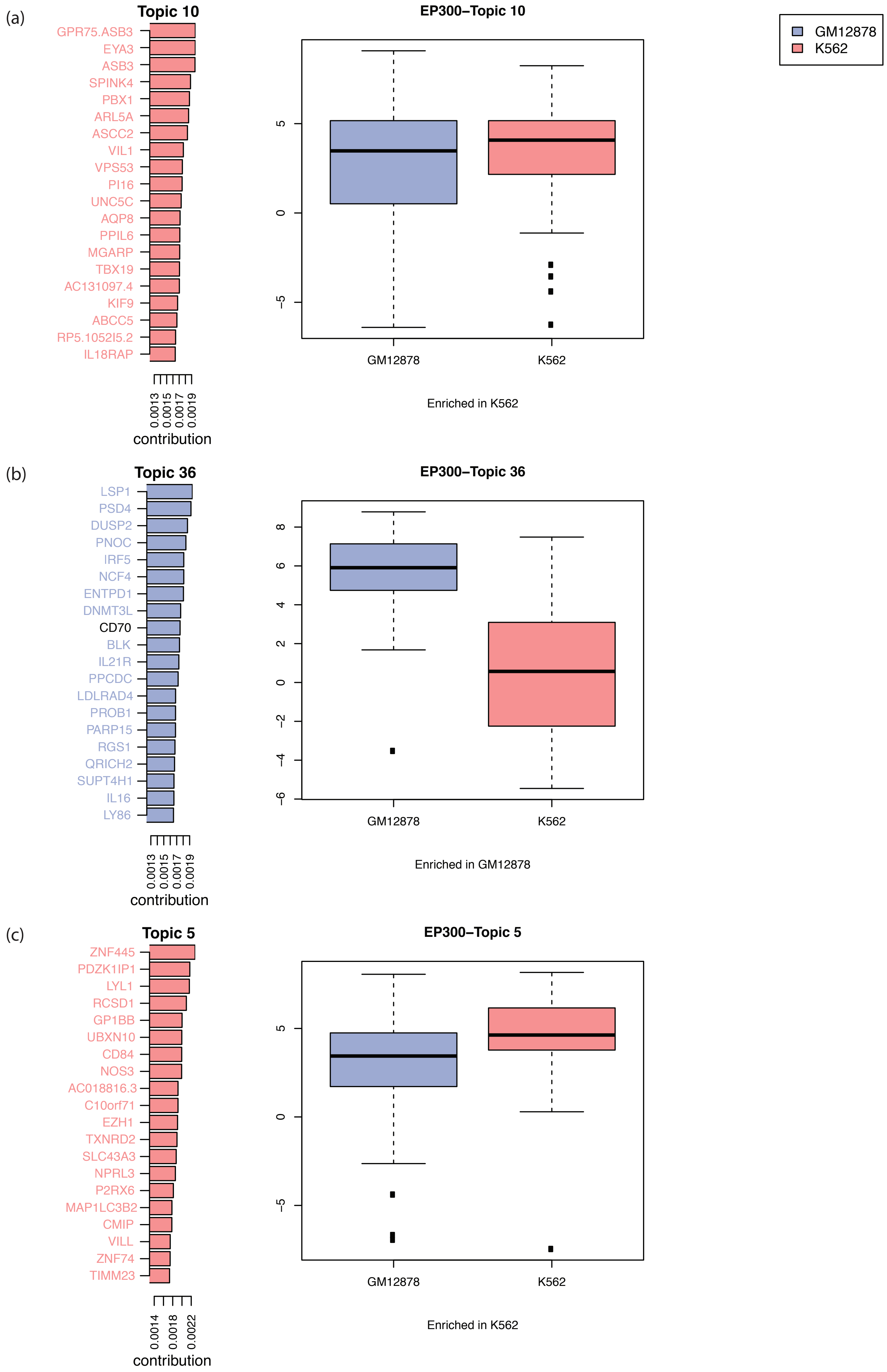
(a)
Topic 10
GPR75.ASB3
EYA3
ASB3
SPINK4
PBX1
ARL5A
ASCC2
VIL1
VPS53
PI16
UNC5C
AQP8
PPIL6
MGARP
TBX19
AC131097.4
KIF9
ABCC5
RP5.1052I5.2
IL18RAP
0.0013
0.0015
0.0017
0.0019
contribution
EP300−Topic 10
5
0
−5
GM12878
K562
Enriched in K562
GM12878
K562
(b)
Topic 36
LSP1
PSD4
DUSP2
PNOC
IRF5
NCF4
ENTPD1
DNMT3L
CD70
BLK
IL21R
PPCDC
LDLRAD4
PROB1
PARP15
RGS1
QRICH2
SUPT4H1
IL16
LY86
0.0013
0.0015
0.0017
0.0019
contribution
EP300−Topic 36
8
6
4
2
0
−2
−4
−6
GM12878
K562
Enriched in GM12878
(c)
Topic 5
ZNF445
PDZK1IP1
LYL1
RCSD1
GP1BB
UBXN10
CD84
NOS3
AC018816.3
C10orf71
EZH1
TXNRD2
SLC43A3
NPRL3
P2RX6
MAP1LC3B2
CMIP
VILL
ZNF74
TIMM23
0.0014
0.0018
0.0022
contribution
EP300−Topic 5
5
0
−5
GM12878
K562
Enriched in K562

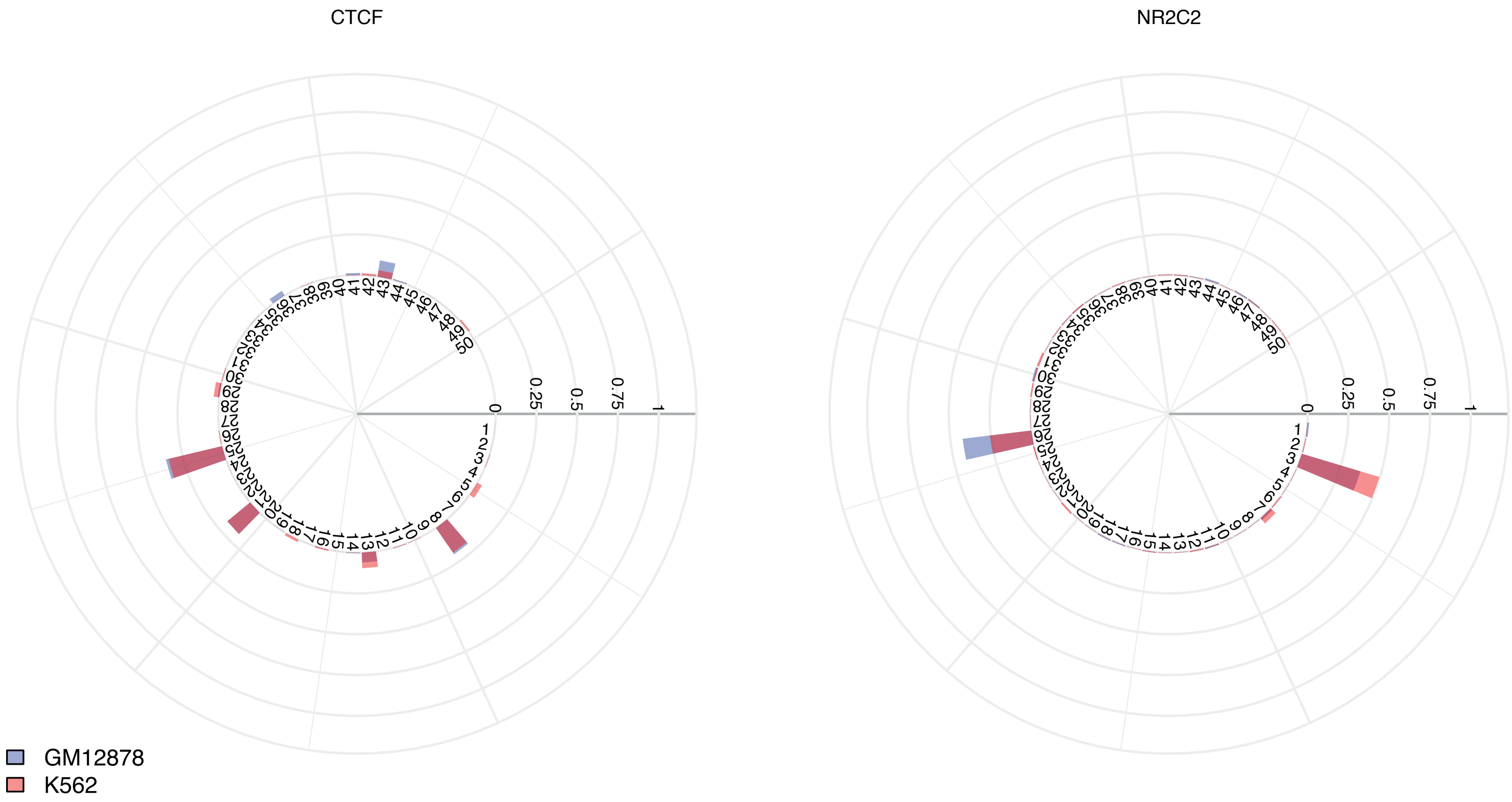

CTCF
NR2C2
0
0.25
0.5
0.75
1
GM12878
K562

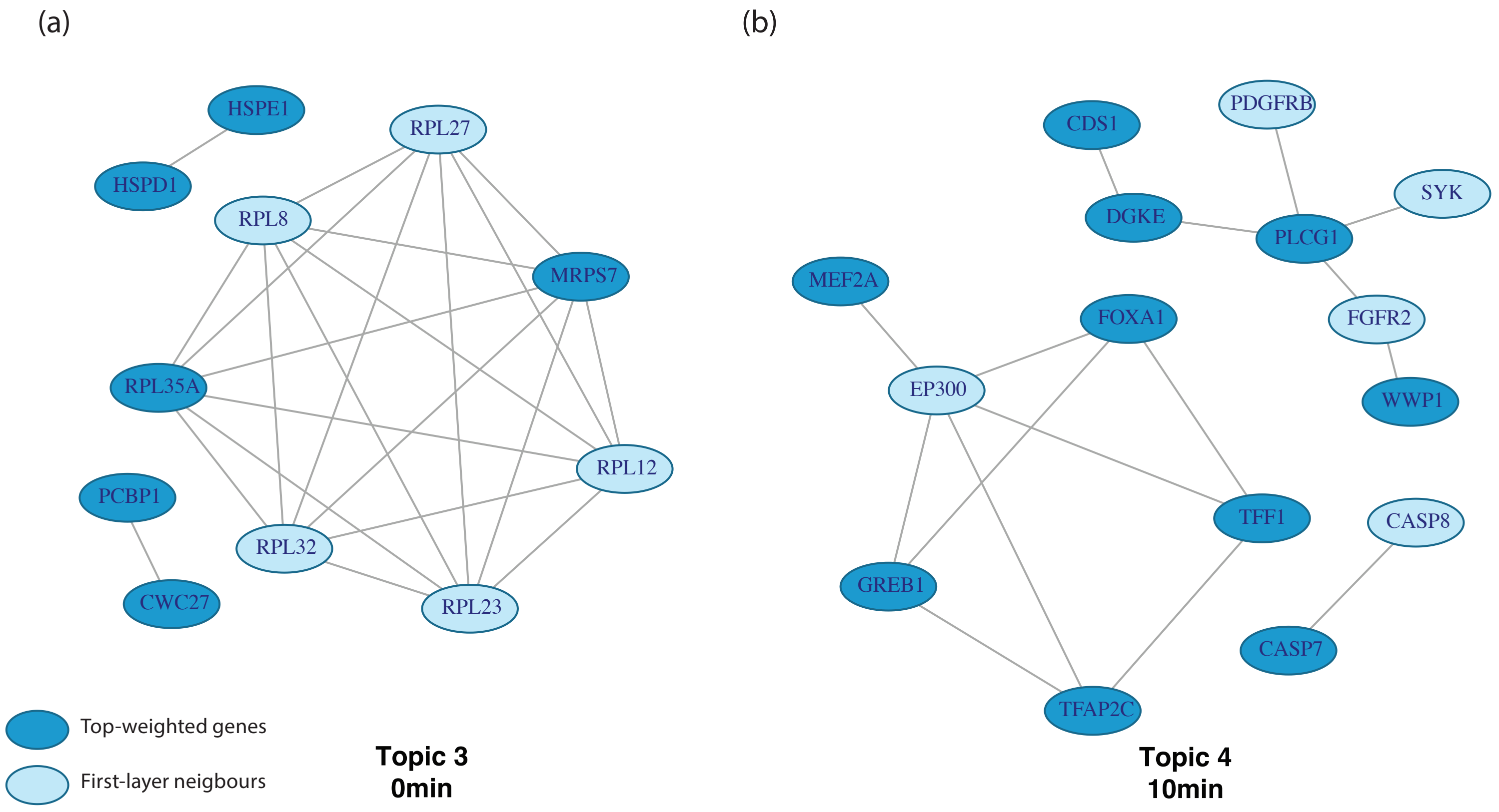
(a)
HSPE1
HSPD1
RPL27
RPL8
MRPS7
RPL35A
RPL12
PCBP1
CWC27
RPL32
RPL23
Top-weighted genes
First-layer neigbours
Topic 3
0min
(b)
CDS1
PDGFRB
SYK
DGKE
PLCG1
MEF2A
FOXA1
FGFR2
EP300
WWP1
TFF1
CASP8
GREB1
CASP7
TFAP2C
Topic 4
10min

(a) Topic 10

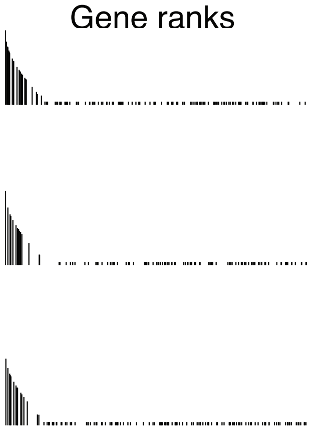

| Pathway | Gene ranks | NES | pval | padj |
|---|---|---|---|---|
| ESC_V6.5_UP_EARLY.V1_DN | | 1.07 | 2.0e−03 | 3.8e−01 |
| CTIP_DN.V1_DN | | 1.06 | 1.9e−02 | 7.2e−01 |
| ATM_DN.V1_UP | | 1.05 | 2.6e−02 | 7.2e−01 |

(b) Topic 26

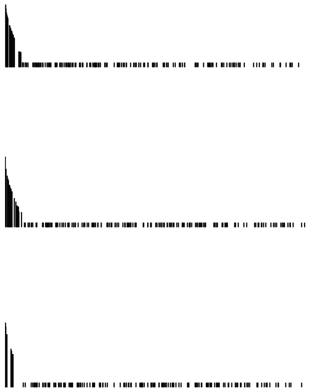

| Pathway | Gene ranks | NES | pval | padj |
|---|---|---|---|---|
| CAMP_UP.V1_UP | | 1.24 | 1.0e−03 | 1.9e−01 |
| RAPA_EARLY_UP.V1_DN | | 1.21 | 3.0e−03 | 2.8e−01 |
| GCNP_SHH_UP_LATE.V1_DN | | 1.22 | 5.0e−03 | 3.1e−01 |

(c) Topic 35

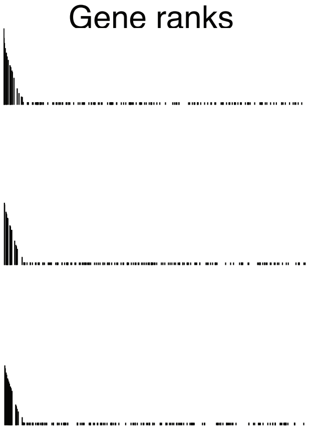

| Pathway | Gene ranks | NES | pval | padj |
|---|---|---|---|---|
| MYC_UP.V1_UP | | 1.07 | 2.0e−03 | 3.8e−01 |
| RPS14_DN.V1_DN | | 1.07 | 4.0e−03 | 3.8e−01 |
| E2F1_UP.V1_UP | | 1.06 | 1.3e−02 | 8.2e−01 |

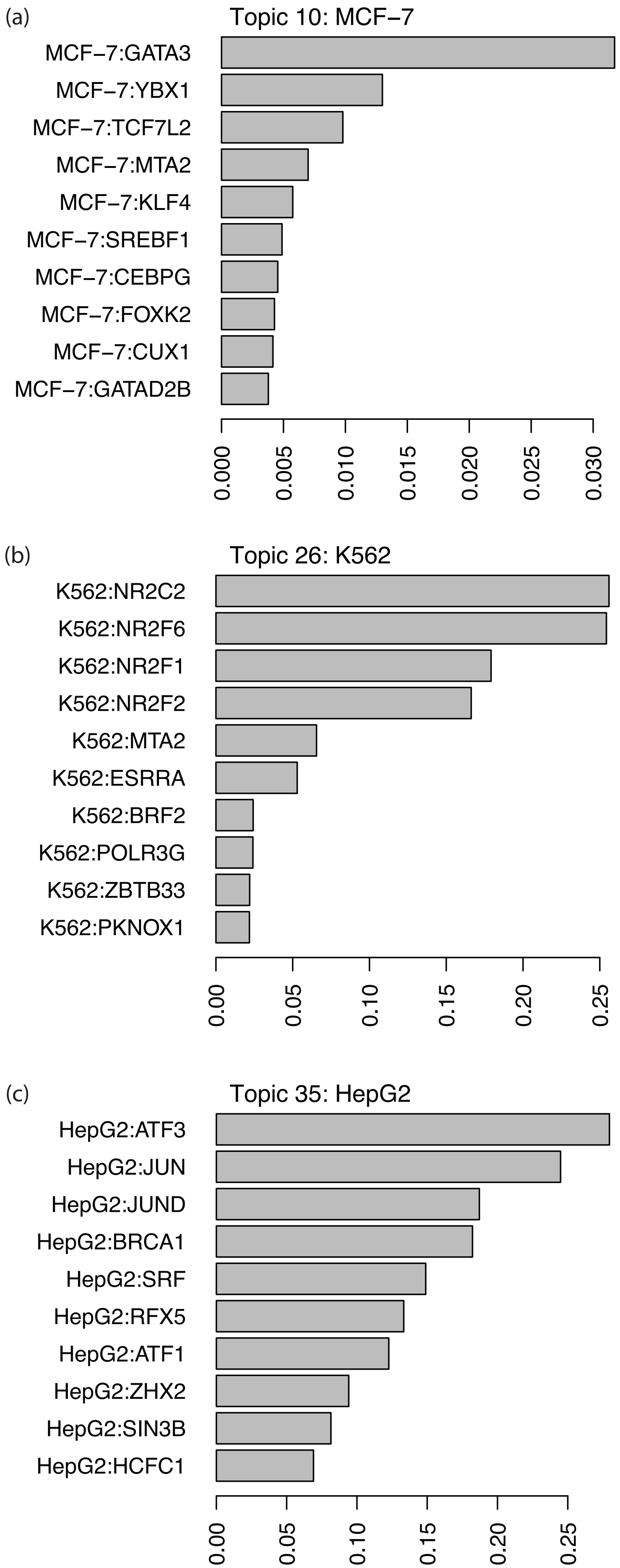

(a)
Topic 10: MCF−7
MCF−7:GATA3
MCF−7:YBX1
MCF−7:TCF7L2
MCF−7:MTA2
MCF−7:KLF4
MCF−7:SREBF1
MCF−7:CEBPG
MCF−7:FOXK2
MCF−7:CUX1
MCF−7:GATAD2B
0.000
0.005
0.010
0.015
0.020
0.025
0.030
(b)
Topic 26: K562
K562:NR2C2
K562:NR2F6
K562:NR2F1
K562:NR2F2
K562:MTA2
K562:ESRRA
K562:BRF2
K562:POLR3G
K562:ZBTB33
K562:PKNOX1
0.00
0.05
0.10
0.15
0.20
0.25
(c)
Topic 35: HepG2
HepG2:ATF3
HepG2:JUN
HepG2:JUND
HepG2:BRCA1
HepG2:SRF
HepG2:RFX5
HepG2:ATF1
HepG2:ZHX2
HepG2:SIN3B
HepG2:HCFC1
0.00
0.05
0.10
0.15
0.20
0.25

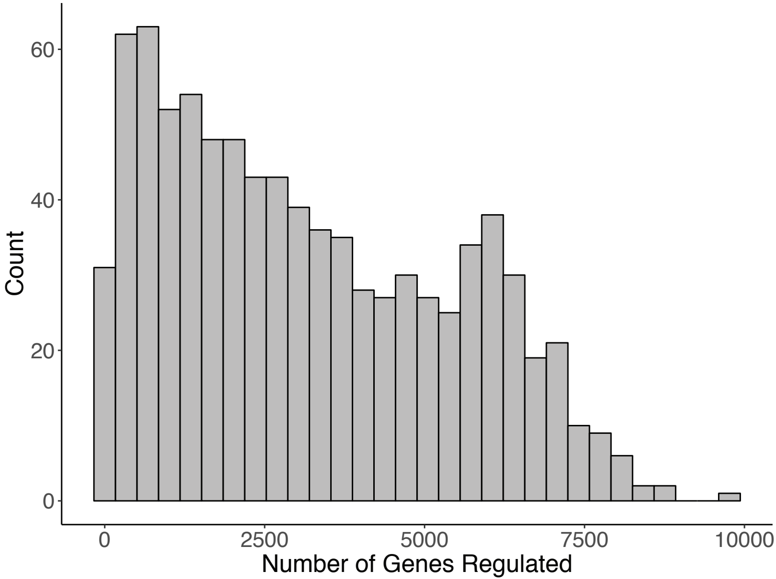

Count
60
40
20
0
0
2500
5000
7500
10000
Number of Genes Regulated

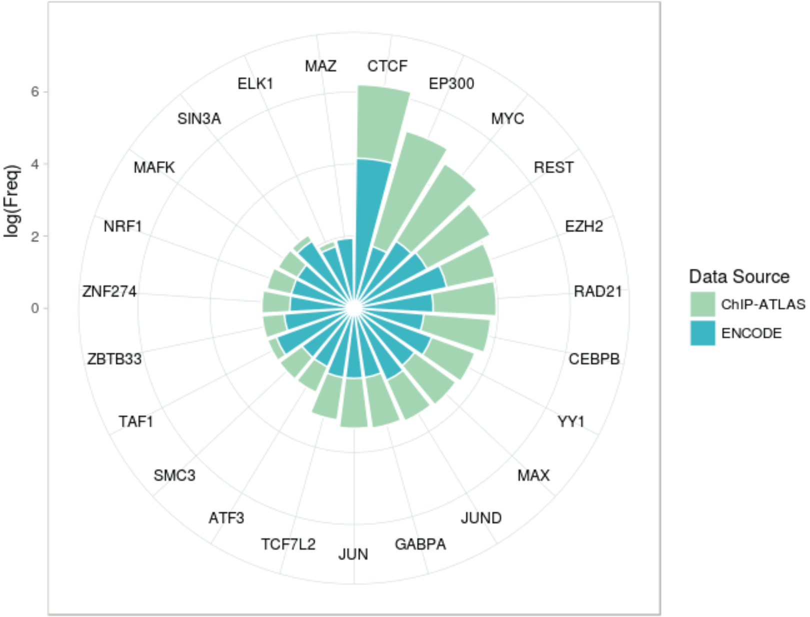

log(Freq)
0
2
4
6
CTCF
EP300
MYC
REST
EZH2
RAD21
CEBPB
YY1
MAX
JUND
GABPA
JUN
TCF7L2
ATF3
SMC3
TAF1
ZBTB33
ZNF274
NRF1
MAFK
SIN3A
ELK1
MAZ
Data Source
ChIP-ATLAS
ENCODE
